# Dynamic Apical-Basal Enrichment of the F-Actin during Cytokinesis in Arabidopsis Cells Embedded in their Tissues

**DOI:** 10.1101/2021.07.07.451432

**Authors:** Alexis Lebecq, Aurélie Fangain, Alice Boussaroque, Marie-Cécile Caillaud

## Abstract

During the life cycle of any multicellular organism, cell division contributes to the proliferation of the cell in the tissues as well as the generation of specialized cells, both necessary to form a functional organism. Therefore, the mechanisms of cell division need to be tightly regulated, as malfunctions in their control can lead to tumor formation or developmental defects. This is particularly true in land plants, where cells cannot relocate and therefore cytokinesis is key for morphogenesis. In the green lineage, cell division is executed in radically different manners than animals, with the appearance of new structures (the preprophase band (PPB), cytokinetic the cell plate and phragmoplast), and the disappearance of ancestral mechanisms (cleavage, centrosomes). While F-actin and microtubules closely co-exist to allow the orientation and the progression of the plant cell division, recent studies mainly focused on the microtubule’s involvement in this key process. Here, we used our recently developed root tracking system to follow actin dynamics in dividing Arabidopsis meristematic root cells. In this study, we imaged in time and space the fluorescent-tagged F-actin reporter Lifeact together with cell division markers in dividing cells embedded in their tissues. In addition to the F-actin accumulation’s in the phragmoplasts, we observed and quantified a dynamic apical-basal enrichment of the F-actin during cytokinesis. The role and the possible actors responsible for F-actin dynamics during cytokinesis are discussed.

## INTRODUCTION

During plant mitosis, the cytoskeleton reorganized in plant-specific structures that allow the separation of the mother cell into two daughter cells by the centrifugal growth of a new wall. While the role of the actin cytoskeleton is of prime importance for animal cell division, the green cells require mostly microtubules in the early step of the cell division. Indeed, the plant cells can divide in the absence of F-actin (Baluska et al. 2001; Nishimura et al. 2003) and division still occurs in the presence of mutations in genes encoding actin or various actin-associated proteins (Jürgens 2005). Before mitosis, the migration of the nucleus is not prevented by the cytochalasin D, a drug that disrupted the actin filaments (F-actin), suggesting that microtubules play an important role in this process (Katsuta et al. 1990). Yet, the actin cytoskeleton is found early during division with the formation of an actin ring at the cortex of the cell in preprophase, the actin preprophase band (PPB) which is considered to be wider than the microtubules PPB (Palevitz 1987).

The formation of actin PPB depends on microtubules because the application of microtubules-depolymerizing drugs prevents the formation of both the microtubules and F-actin components of the PPB (Palevitz 1987). However, actin PPB can also affect the microtubules PPB because actin depolymerization results in drastic enlargement of the microtubules PPB and fluctuating division planes (Mineyuki and Palevitz 1990). As the microtubules PPB narrows, the actin PPB narrows, and when the microtubules PPB reach their narrowest configuration, the fluorescence signals of F-actin start to disappear from the cell cortex region occupied by the microtubules PPB, giving rise to the actin-depleted zone (Cleary et al. 1992; Liu and Palevitz 1992). In tobacco BY-2 cell culture cells, this actin-depleted zone is unambiguously visible and flanked by enrichments of F-actin while this depleted zone was not yet visualized in plant cells embedded in their tissues. Whether the actin-depleted zone coincides with the cortical division zone which acts as a landmark for the future cell division site is still unclear. Nevertheless, the actin-depleted zone is short-lived from metaphase through the anaphase/telophase transition (Kakimoto and Shibaoka 1987; Clayton and Lloyd 1985).

While cytokinesis is progressing, the phragmoplast that guides the delivery of vesicles to form the expanding cell plate is composed of both microtubules and actin filaments (Staehelin and Hepler 1996). Live-cell imaging over time in BY2 cells revealed that F-actin was involved in endoplasmic reticulum accumulation in the phragmoplast at the late phase (Higaki et al. 2008). Tobacco BY-2 cells treated with F-actin-depolymerizing drugs show disorganized phragmoplasts and wrinkled cell plates (Hoshino et al. 2003; Yoneda et al. 2004; Sano et al. 2005; Higaki et al. 2008; Kojo et al. 2013), which uncover the role of the F-actin during plant cytokinesis.

Because F-actin and microtubules closely co-exist and play important roles in the phragmoplast (Smith 1999), cooperation or interaction between of actin and microtubules cytoskeletons is assumed, and several proteins have been proposed to mediate the cooperation or interaction between actin and microtubules cytoskeletons (Igarashi et al. 2000; Buschmann et al. 2011; Klotz and Nick 2012; Schwab et al. 2003; Maeda et al. 2020). In particular, the moss myosin VIII, a motor protein that interacts with both actin and microtubules, localizes to the division site and the phragmoplast, linking microtubules to the cortical division zone via actin cytoskeleton (Wu and Bezanilla 2014). These findings, together with other reports in which drug treatments and localization studies were performed (Lloyd and Traas 1988; Molchan et al. 2002; Takeuchi et al. 2016), highlight that actin cytoskeleton may interact with the microtubules bridging the cell cortex and the phragmoplast during cytokinesis.

Early work using immunolocalization and gold immunolabeling of the actin cytoskeleton revealed the presence of a meshwork of F-actin that concentrated at the cortex of the cell, on the side facing the spindle poles (Mole-Bajer et al. 1988; Cho and Wick 1991). Recently, super-resolved imaging of the F-actin was performed using a photo-activatable Lifeact, a 17-amino-acid peptide that stained F-actin structures in BY2 cells (Durst et al. 2014). Using this approach, a gradient of the Lifeact–psRFP signal was observed that progressively led to a clear increase of signal intensity toward the cell poles, especially during the later mitotic phases (Durst et al. 2014). However, the dynamic of such polarization was not yet reported in dividing plant cells surrounded by its tissue.

Here, we used our recently developed root tracking system to follow actin dynamics in dividing Arabidopsis meristematic root cells. By the visualization of the fluorescent-tagged Lifeact together with cell division markers, we observed an apical-basal accumulation of the F-actin in the late stage of cell division in Arabidopsis root meristematic cells. The role and the possible actors responsible for this actin enrichment are discussed.

## RESULTS

In order to analyze in vivo the behaviors of the actin cytoskeleton in a dividing tissue, we collected a set of Lifeact fluorescent reporters, tagged with YFPv (Doumane et al. 2021), tdTomato (Liao and Weijers 2018). We generated a blue-tagged version of Lifeact with 2xmTURQUOISE2 under a Ub10 promoter (Lifeact-2xmTU2). Using the recently developed root tracking system, we observed the behavior of the actin cytoskeleton every three minutes in epidermal cells of Arabidopsis up to five root tips in one experiment, in three dimensions, using a spinning disk confocal microscope (Doumane et al. 2017). We first analyze in time-lapse Arabidopsis meristematic root cells during division in plants expressing the actin marker *Lifeact-YFPv* (Figure 1). The z-projection of division tissues expressing the actin marker Lifeact-YFPv showed that during cytokinesis, the F-actin is clearly observed in the developing phragmoplast (Figure 1, empty arrows). Moreover, an apical-basal accumulation of Lifeact-YFPv during cytokinesis was clearly visible in the dividing root tissues (Figure 1, white plain arrows).

**Figure 1.**
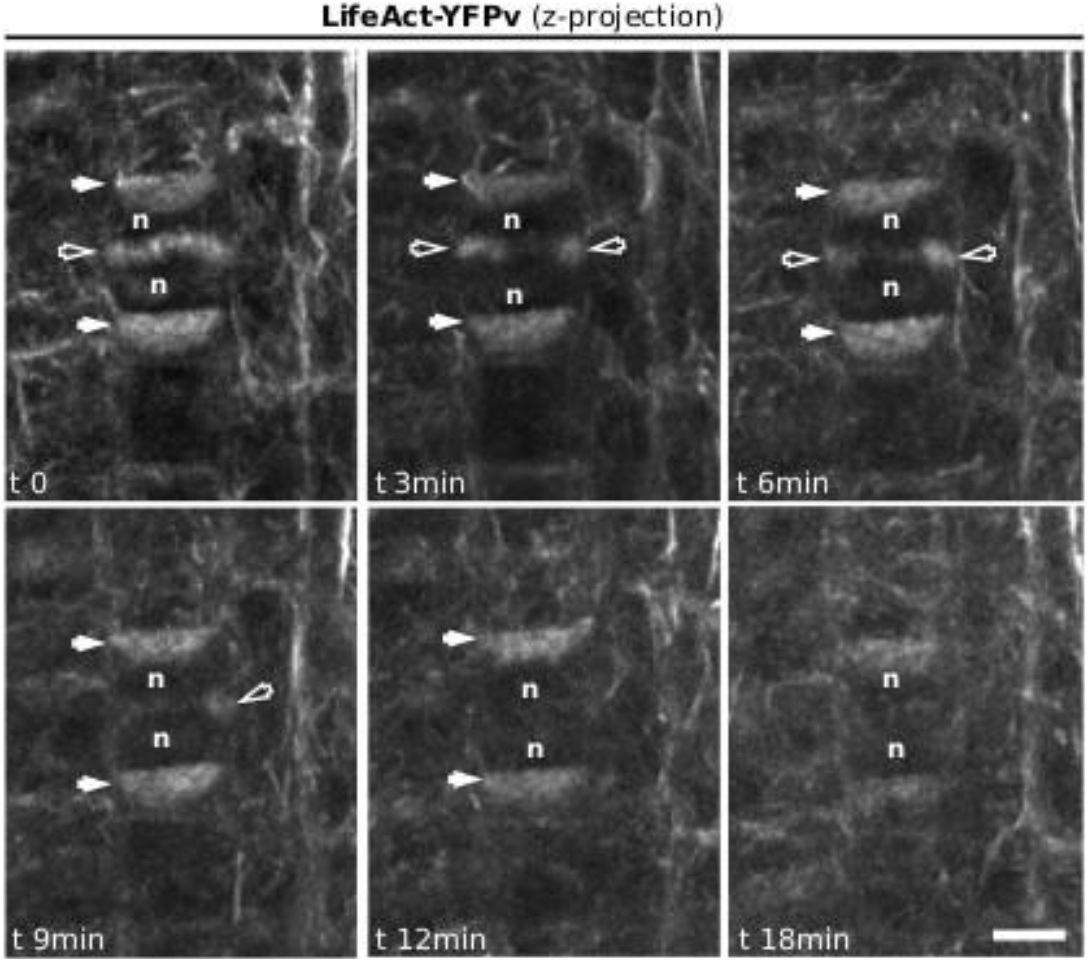
Representatives images of a z-projection of a time-lapses analysis in Arabidopsis root meristem, expressing *Ub10:Lifeact-YFPv*. White arrows: F-actin enrichment at the apical-basal part of the cells; empty arrows: F-actin in the phragmoplast; n: nucleus. Scale bar = 5 μm.

We confirmed the F-actin patterning during cell division obtained with the LifeAct-YFPv reporter with another well characterized F-actin biosensor. We performed the same analysis with plant expressing the C-terminal half of the plant Fimbrin-like gene AtFim1 (aa 325–687) fused in both sides to the GFP fluorescent protein (m*GFP-ABD2)*. Time-lapses analysis of the growing root Arabidopsis meristem expressing m*GFP-ABD2* displayed the same apical-basal accumulation of F-actin during cytokinesis (Figure 2).

**Figure 2.**
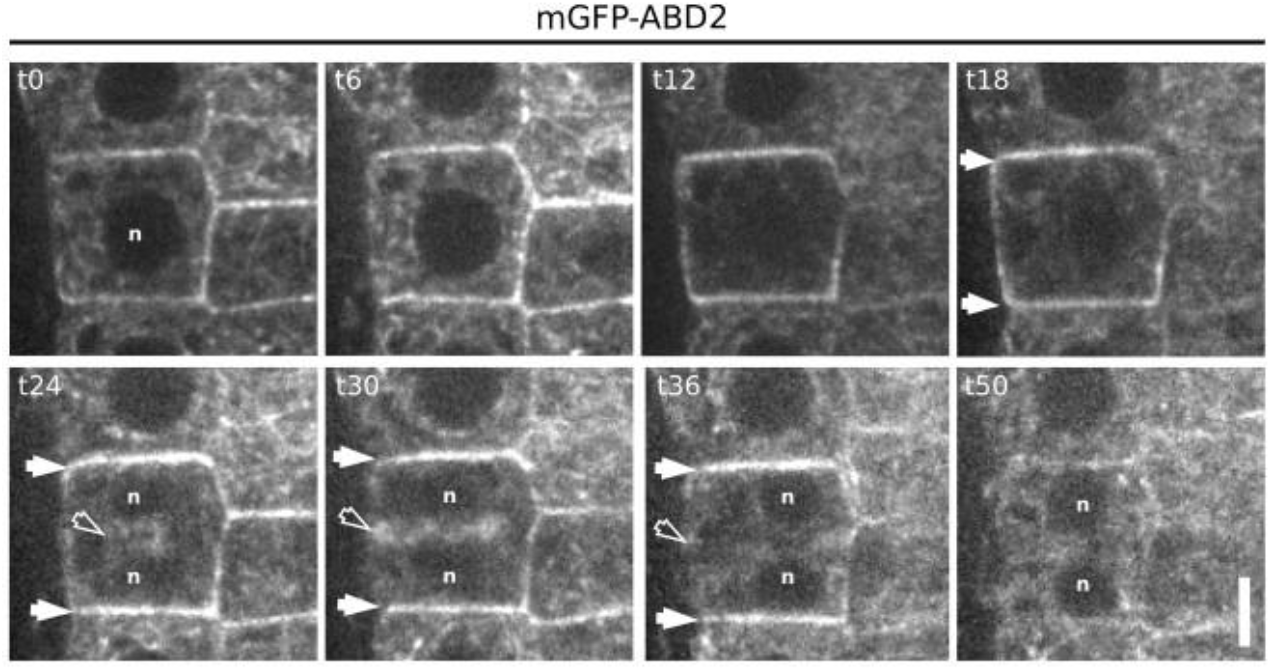
Representatives images of a z-projection of a time-lapses analysis in Arabidopsis root meristem, expressing *mGFP-ABD2*. White arrows: F-actin enrichment at the apical-basal part of the cells; empty arrows: F-actin in the phragmoplast; n: nucleus. Scale bar = 5 μm.

We projected in the z-plane the root cells expressing and by changing the orientation to enable the observation of the apical-basal face of the dividing cell (Figure 3A). Using this approach, we confirmed that while the phragmoplast was extending, the Lifeact-2xmTU2 signal covered the entire apical-basal plane of the cell while no accumulation was observed in the lateral part of the cell (Figure 2A). During phragmoplast expansion, the signal at the apical-basal part of the cell stayed strong before the decrease in the fluorescence signal could be observed at the end of the cytokinesis (Figure 1–3).

**Figure 3.**
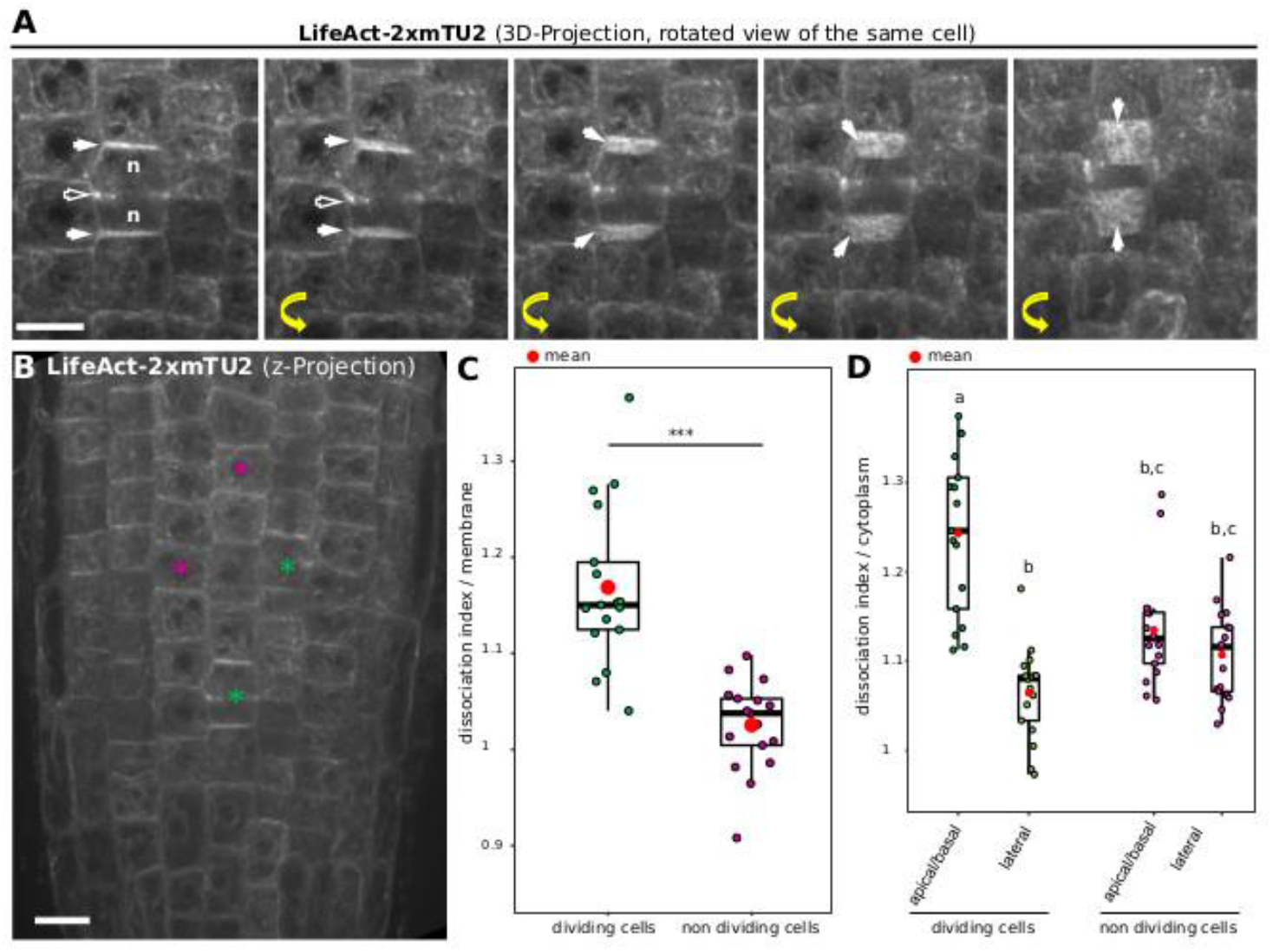
A) Representative images of a three dimensional projection of a dividing cell in Arabidopsis root meristem expressing *Ub10:LifeAct-2xmTU2*. White arrows, F-actin enrichment at the apical-basal part of the cells; empty arrows, F-actin in the phragmoplast; n: nucleus. The yellow arrows marked the four rotated images obtained by the rotation in imageJ of the initial z-projected image presented in the left panel. Scale bar = 10 μm. B) representative images of the distribution of LifeAct-2xmTU2 in dividing cell and non-dividing cells used for the quantification. Scale bar = 15 μm. C) Quantification of the localization index of LifeAct-2xmTU2 between the apical-basal membranes and the lateral membranes of dividing vs non-dividing cells. D) Quantification of the localization index of LifeAct-2xmTU2 between the apical-basal membrane or the lateral membranes and the cytoplasm of dividing vs non-dividing cells. In the plots, the middle horizontal bars represent the median, while the bottom and top of each box represent the 25th and 75th percentiles, respectively. At most, the whiskers extend to 1.5 times the IQR and exclude data beyond that range. For the range of values under 1.5 times the IQR, the whiskers represent the range of maximum and minimum values.

We quantify the observed signal in the Arabidopsis line expressing *LifeAct-2xmTU2* in dividing cell and non-dividing cells. To do so, we analyze using ImageJ the ratio of fluorescence intensity between the two apical and basal membranes and the fluorescence intensity in the lateral membranes (Figure 3B-C). We analyzed the ratio of fluorescence intensity between the apical-basal membrane and the cytoplasm for dividing cell and non-dividing cells (Figure 3B-D). Using this quantitative approach, we confirmed the enrichment of the F-actin signal at the apical/basal part of the dividing cells, while no specific distribution was observed in non-dividing cells.

To address whether the actin depleted zone was quantifiable, we analyze the ratio of fluorescence intensity between the lateral membranes and the cytoplasm for dividing cell and non-dividing cells (Figure 3B-D). While a slight decrease in the ratio is observed for the dividing cells compared to non-dividing cells, the presence of the adjacent non-dividing cells surrounding this region might mask a clear depletion. Indeed, while the actin depleted zone is clearly observed in cell culture (Cleary et al. 1992; Liu and Palevitz 1992), this depleted zone is difficult to image in dividing tissue (Voigt et al. 2005).

Using plant co-expressing *Lifeact-tdTom* and the cell plate marker *YFP-KNOLLE*, we tracked dividing cells in the root meristem from early metaphase. While the spindle was forming, and F-actin was observed at the spindle, the polarization of the actin cytoskeleton in the apical-basal part of the cell was not yet observed (Figure 4). An increase of the signal could be observed at the periphery of the cell, flanking the cortical division zone, but the signal in the surrounding cell was masking the signal from the dividing cells (Figure 4, white arrows), therefore it did not allow us to certify that this signal corresponds to the earlier described “actin twin peaks” (Smertenko et al. 2010). But as soon as YFP-KNOLLE was found enriched in the developing cell plate, the lateral signal corresponding to Lifeact-tdTom disappeared and the rapid accumulation of Lifeact-tdTom signal at the apical-basal part of the cell was observed, as well as in the phragmoplast (Figure 4). This accumulation of F-actin at the apical-basal part of the dividing cell was stable during cell plate expansion as YFP-KNOLLE signal expanded toward the cell periphery (Figure 4). When the cell plate was fully attached to the mother cell, no more accumulation of the F-actin at the pole of the cell could be observed using the Lifeact-tdTom reporter line (Figure 4). At this stage, the cable of F-actin was found in the cytoplasm of newly formed daughter cells (Figure 4).

**Figure 4.**
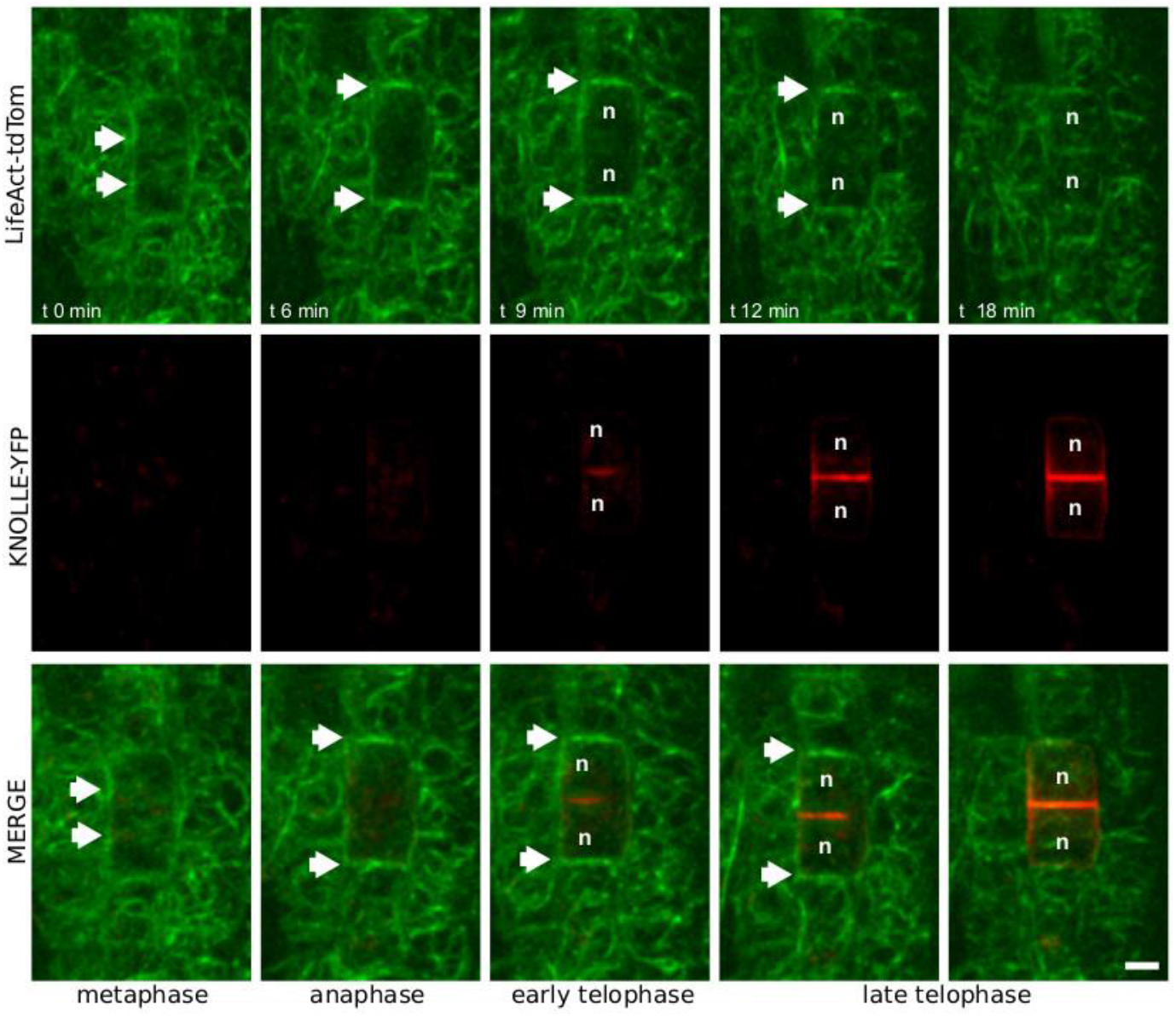
Representatives images of a time-lapses analysis in Arabidopsis root meristem, expressing RPS5a:LifeAct-tdTom (green) together with KNOLLE-YFP (red). White arrows, F-actin enrichment at the apical-basal part of the cells; n: nucleus. Scale bar = 5 μm.

We confirmed these results using an Arabidopsis line co-expressing the cortical division zone marker GFP–PHGAP1 together with Lifeact-tdTom (Figure 6). In transgenic lines expressing *GFP–PHGAP1* under a *Ub10 promoter*, the fluorescent signal was reported to accumulate at the cortical division zone in meta-/anaphase and remains throughout cytokinesis (Stockle et al. 2016). Dual localization of GFP-PHGAP1 and Lifeact-tdTom showed that the accumulation at the apical/basal part of the dividing cells appeared before the signal corresponding to GFP–PHGAP1 could be noticed, suggesting that the F-actin polarization appears before the recruitment of PHGAP1 at the CDZ (Figure 6, supplemental movie 1).

**Figure 5.**
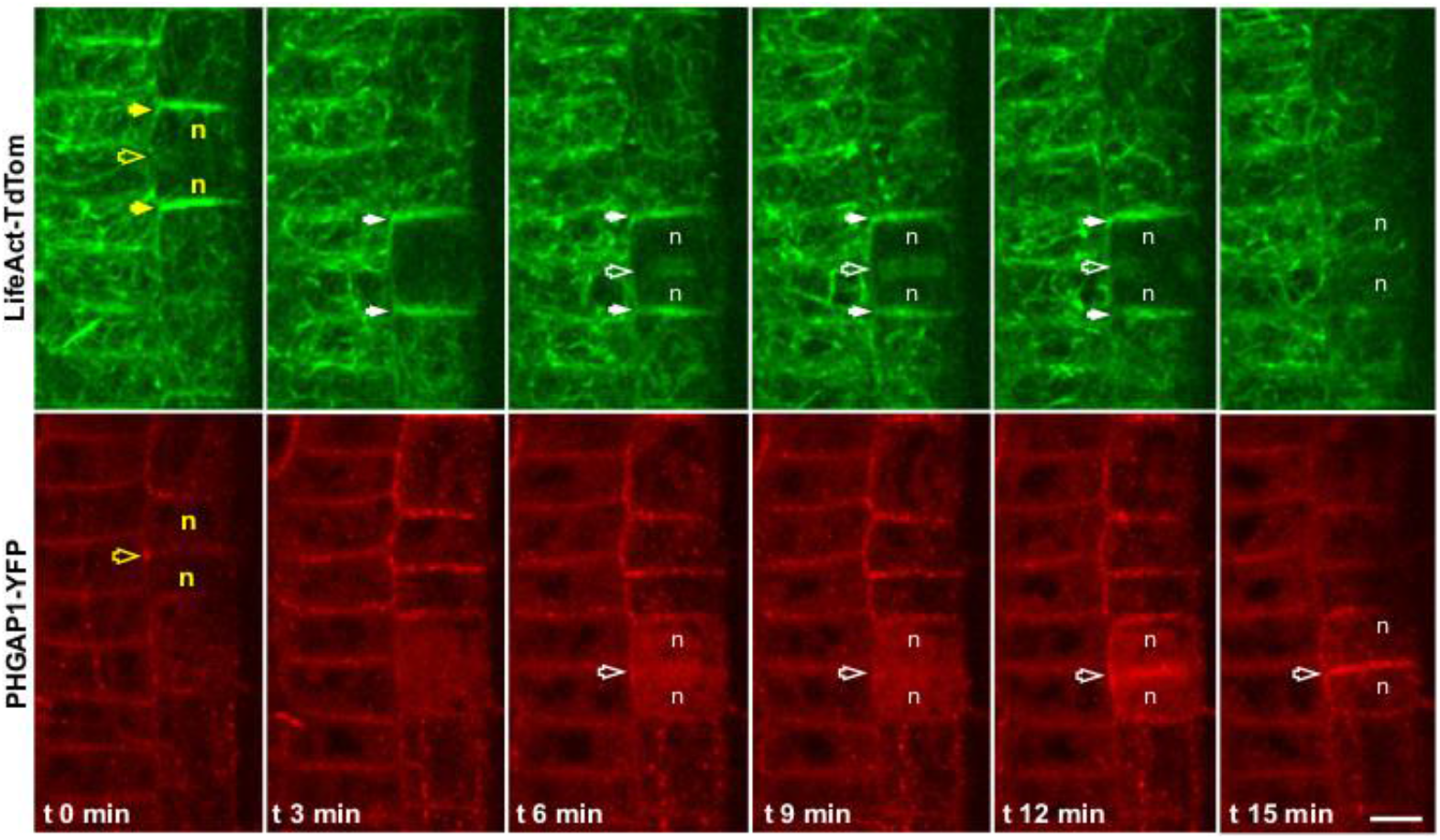
Representatives images of a time-lapses analysis in Arabidopsis root meristem, expressing RPS5a:LifeAct-tdTom (green) together with PHGAP1-YFP (red). Yellow and white arrows, F-actin enrichment at the apical-basal part of the cells; empty arrows, position of the cortical division zone; n: nucleus. Scale bar = 5 μm.

**Figure 6.**
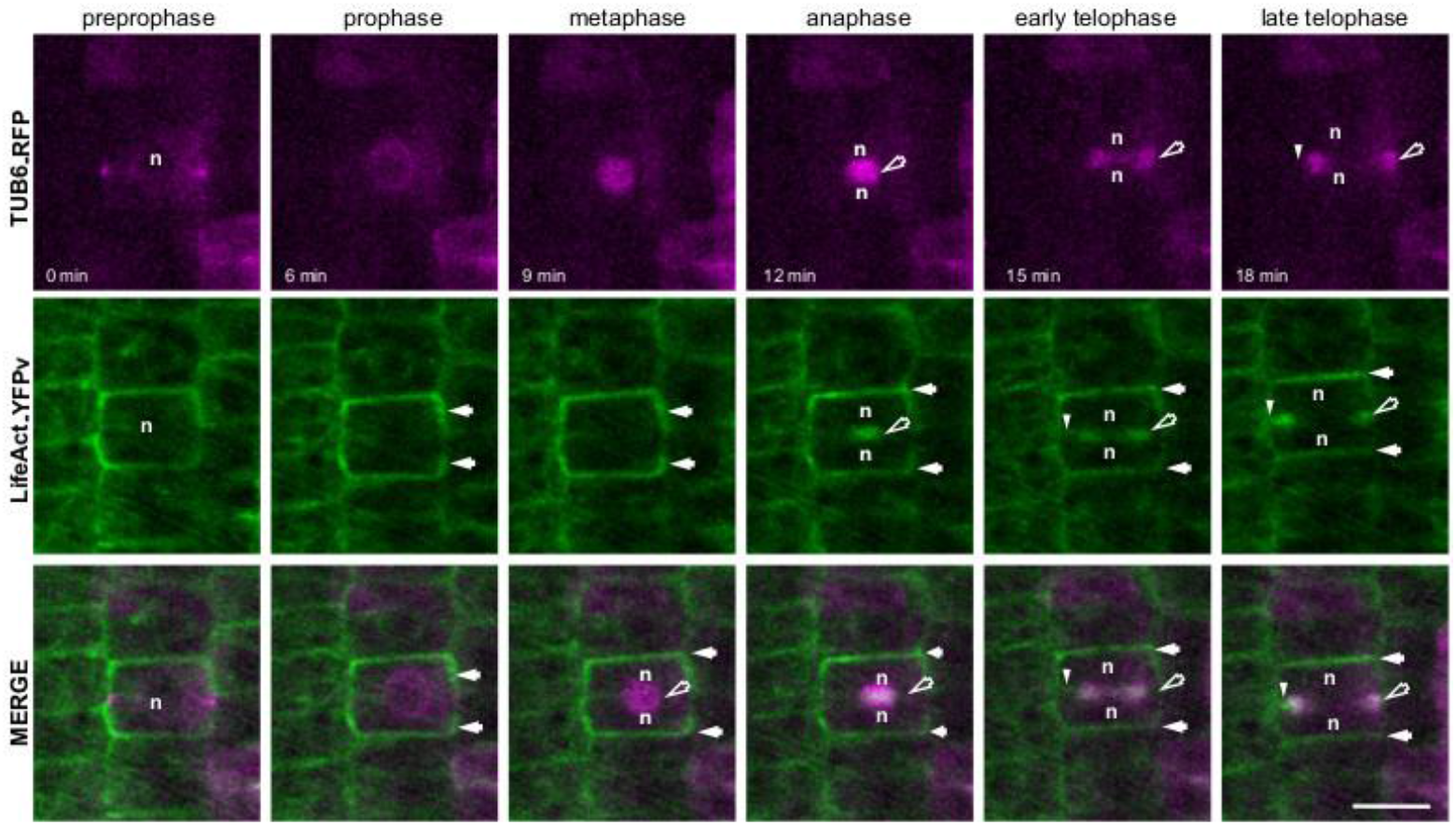
Representatives images of a time-lapses analysis in Arabidopsis root meristem, expressing Ub10:LifeAct-YFPv (green) together with TUA6-RFP (magenta). White arrows, F-actin enrichment; empty arrows, the phragmoplast; arrow head, position of the F-actin in the phragmoplast edge; n: nucleus. Scale bar = 10 μm.

Colocalization of F-actin using Lifeact-tdTom with mitotic microtubules decorated by TUA6-RFP allowed used to visualize in detail the dynamics at the phragmoplast. During early anaphase, the colocalization of microtubules and F-actin was total while in late cytokinesis, colocalization analysis highlighted the differences in positioning of both cytoskeletons (Figure 6, supplemental movie 1). At the ring phragmoplast stage, the lifeact-tdTom preceded the TUA6-RFP at the edge of the phragmoplast (Figure 6, supplemental movie 1), confirming the role of F-actin in the phragmoplast guidance in the attachment phases of the cytokinesis.

## DISCUSSION

An extraordinary new view of the dynamic spatiotemporal regulation of keys components of the cell division orientation is emerging from advances in live-cell imaging in Arabidopsis root meristem (Muller et al. 2006; Walker et al. 2007; Xu et al. 2008; Drevensek et al. 2012; Lipka et al. 2014; Stockle et al. 2016; Schaefer et al. 2017). However, when it comes to studying subcellular dynamics during plant cell division manual adjustment is required since the area under study will rapidly grow out of the field of view. This makes it necessary to obtain an automated system to release the biology researcher from spending up to a day in front of the microscope performing adjustments every ten minutes. In the present study, we took advantage of a confocal chasing system to track the meristematic zone of the root based on point-to-point correspondences and motion estimation (Doumane et al. 2017). Using this approach, we can track over a long period (10 hours) mitosis in the meristem of multiple Arabidopsis roots by spinning disk confocal microscopy. This approach allowed us to rapidly collect biological replicates on the behavior of the actin cytoskeleton during plant cell division and to quantify it. We showed that, in addition to its localization in the phragmoplast, F-actin polarized at the apical-basal part of the cell from the meta-anaphase transition and decreased at the end of the cytokinesis to eventually formed again cables in the cytoplasm of the two newly formed daughter cells. How this transient accumulation is achieved remains unclear.

In animal cytokinesis, the local increase in the level of the anionic lipid PI(4,5)P_2_ modifies F-actin amounts at the cell equator and controls cytokinesis abscission (Dambournet et al. 2011). Using our root tracking system, we observed that PI(4,5)P_2_ has specific subcellular localization during plant cell division (Simon et al. 2016) suggesting a link between the proper spatial production of phosphoinositides and successful completion of cytokinesis. In particular, we found that PI(4,5)P_2_ also seems to polarly localized during cell division (Simon et al. 2016) with a patterning resembling the one observed for F-actin in this study. Moreover, depletion of the PI(4,5)P_2_ from the plasma membrane using the iDePP system highlighted the importance of P I(4,5)P_2_ pool for proper F-actin dynamics (Doumane et al. 2021). Actin nucleators such as ARP2/3 were shown to interact strongly with the plasma membrane in Arabidopsis (Kotchoni et al. 2009), and both ACTIN DEPOLYMERIZATION FACTOR (ADF)/cofilin and profilin bind to PI(4,5)P_2_ (Gungabissoon et al. 1998). In *Zea mays*, PROFILIN1 (ZmPRO1) inhibits hydrolysis of PI(4,5)P_2_ at the membrane by the membrane-associated enzyme Phospholipase C (*PLC*)(Staiger et al. 1993). It was reported that PROFILIN from *Phaseolus vulgaris* could bind directly to the phosphoinositide enzyme PI-3Kinase, which suggests that PROFILIN might participate in membrane trafficking, and acts as a linker between the endocytic pathway and the F-actin dynamics (Aparicio-Fabre et al. 2006).

In Arabidopsis, enzymes that locally produce PI(4,5)P_2_ named PIP5K1 and PIP5K2 were reported to be most active among the ubiquitously expressed PI4P 5-kinases (Ischebeck et al. 2008) and therefore are likely making a major contribution to cellular PI(4,5)P_2_ formation (Tejos et al. 2014). During embryogenesis, double mutations in *pi5k1/pi5k2* bent to defects characterized by the formation of a vague apical-to-basal boundary (Tejos et al. 2014) which eventually lead to sterile plants (Tejos et al. 2014). Functional translational fusions with a yellow fluorescent protein (YFP) were generated to visualize the subcellular localization of PIP5K1 and PIP5K2 (Ischebeck et al. 2013). In this study, both YFP-PIP5K1 and YFP-PIP5K2 seems to be enriched in the apical-basal membrane in dividing cells as observed for the F-actin in the present study. However, their dynamic localization during cell division needs yet to be addressed in details, to disentangle if those kinases locally increasing PI(4,5)P_2_ production. Recently developed tools to increase or decrease the pool of PI(4,5)P_2_ at the plasma membrane (Gujas et al. 2017; Doumane et al. 2021) might help us to address such a question in the future. Based on this observation, it is appealing to emphasize that the composition in phosphoinositides could mediate the F-actin polymerization at the apical-basal membranes, allowing the polarization of the cell during its division. Why cells in their tissues needs this F-actin accumulation? It is striking to observe, when it comes to cell division orientation, that in mutants impaired in cortical dividing zone (CDZ) definition and/or microtubules PPB organization, the defects rely on titled cell plates, yet the attachment is still happening in the lateral membranes of the root meristematic cells (Kumari et al. 2021; Schaefer et al. 2017; Stockle et al. 2016; Muller et al. 2006). In other word, in microtubule defective mutants impaired in PPB or CDZ formation, we do not observe the attachment of the cell plate in the apical basal membranes. Challenging the F-actin accumulation at the apical-basal part of the cell might be the key to understand how the cell polarity is maintained after microtubules cytoskeleton perturbation during cell division in plant tissues.

## MATERIAL AND METHODS

### Growth condition and plant materials

*Arabidopsis* Columbia-0 (Col-0) accession was used as a reference genomic background throughout this study. *Arabidopsis* seedlings *in vitro* on half Murashige and Skoog (½ MS) basal medium supplemented with 0.8% plant agar (pH 5.7) in continuous light conditions at 21 °C. Plants were grown in soil under long-day conditions at 21 °C and 70% humidity 16 h daylight.

### Cloning

The lifeact sequence was flanked with attB2R and attB3 sequences and recombined by BP gateway reaction into *pDONR-P_2_RP3*. Final destination vectors were obtained using three fragments LR recombination system (life technologies, www.lifetechnologies.com/) using pB7m34GW destination vector (Karimi et al. 2007).

### Plant material

| construct | ecotype | reference | resistance |
| --- | --- | --- | --- |
| Ub10pro:Lifeact-YFPv | Col-0 | Doumane et al., 2021 | Basta |
| RPS5apro:Lifeact-tdTomato | Col-0 | Liao et al., 2018 | Basta |
| Ub10pro:Lifeact-2mTU2 | Col-0 | this study | Kana |
| KNOLLEpro:YFP-KNOLLE | Ler/ Niederzenz-0 background | (El Kasmi *et al.* , 2013), | Basta |
| pUB10:GFP-PHGAP1 | Col-0 | (Stockle et al. 2016) | Basta |
| TUB6-RFP | Col-0 | (Stockle et al. 2016) | Hygro |
| mGFP-ABD2 | Col-0 | N799991 | Basta |

### Spinning-disk microscopy imaging

Images of the time-lapse images were acquired with the following spinning disk confocal microscope set up: inverted Zeiss microscope (AxioObserver Z1, Carl Zeiss Group, http://www.zeiss.com/) equipped with a spinning disk module (CSU-W1-T3, Yokogawa, www.yokogawa.com) and a ProEM+ 1024B camera (Princeton Instrument, http://www.princetoninstruments.com; Figure 2B only) or Camera Prime 95B (www.photometrics.com) using a 63× Plan-Apochromat objective (numerical aperture 1.4, oil immersion). GFP and mCITRINE were excited with a 488 nm laser (150 mW) and fluorescence emission was filtered by a 525/50 nm BrightLine! a single-band bandpass filter (Semrock, http://www.semrock.com/), tdTom was excited with a 561 nm laser (80 mW) and fluorescence emission was filtered by a 609/54 nm BrightLine! a single-band bandpass filter (Semrock, http://www.semrock.com/). For quantitative imaging, pictures of epidermal root meristem cells were taken with detector settings optimized for low background and no pixel saturation. Care was taken to use similar confocal settings when comparing fluorescence intensity or for quantification. Root tracking experiments were performed as described in (Doumane et al. 2017).

### Statistical analysis

For localization index analysis, one dividing cell and non-dividing cell were taken for each root (17 roots). For each two cells, the signal intensity was measure for the Apical-basal membranes, the two laterals membranes and in two elliptical regions of interest (ROIs) of the cytosol. Ratios were obtained by taking the average of the two measured intensities. Localization index comparison were performed using a Kruskal-Wallis rank sum test analyses in R (v. 3.6.1, (R Core Team, 2019), using R studio interface and the packages ggplot2 (Wickham, 2016). Graphs were obtained with R and R-studio software and customized with Inkscape (https://inkscape.org).

| Kruskal-Wallis |  |  |  |  |
| --- | --- | --- | --- | --- |
| Model | Pair | $\chi^2$ | df | P-value |
| Actin localization index per membrane | dividing cell<br>non-dividing cells | 20.828 | 1 | 5.024e-06 |

| Kruskal-Wallis |  |  |  | Dunn test, method "Bonferroni" |  |
| --- | --- | --- | --- | --- | --- |
| Model | $\chi^2$ | df | P-value | Pair | P-value |
| Actin localization index per cytosol | 20.828 | 1 | 5.024e-06 | Apical-basal membranes of dividing cells /cytosol<br>lateral membranes of dividing cells /cytosol | 0.0000* |
|  |  |  |  | Apical-basal membranes of dividing cells /cytosol<br>Apical-basal membranes of non-dividing cells /cytosol | 0.0174* |
|  |  |  |  | Apical-basal membranes of dividing cells /cytosol<br>lateral membranes of non-dividing cells /cytosol | 0.0004* |
|  |  |  |  | lateral membranes of dividing cells /cytosol<br>Apical-basal membranes of non-dividing cells /cytosol | 0.0149* |
|  |  |  |  | lateral membranes of dividing cells /cytosol<br>lateral membranes of non-dividing cells /cytosol | 0.2305 |
|  |  |  |  | Apical-basal membranes of non-dividing cells /cytosol<br>lateral membranes of non-dividing cells /cytosol | 0.8940 |

## ACKNOWLEDGMENTS

We are grateful to the SiCE group (RDP, Lyon, France) in particular Y. Jaillais, V. Bayle and C. Miège for comments and discussions. We thank Y. Boutté (LBM, Bordeaux, France), for sharing the KNOLLE-YFP marker, Sabine Mueller (University of Tübingen, Germany) for sharing the PHGAP1-YFP reporter and Dolf Weijers (Wageningen University, Netherlands) for sharing the LifeAct-tdTom. We thank P. Bolland, and A. Lacroix from our plant facility. We acknowledge the contribution of SFR Biosciences (UMS3444/CNRS, US8/Inserm, ENS de Lyon, UCBL) facilities: C. Lionet, E. Chatre and J. Brocard at the LBI-PLATIM-MICROSCOPY for assistance with imaging. This work was supported by the French National Research Agency (INTERPLAY; ANR-16-CE13-0021; MCC, AF) and PLANTSCAPE (ANR-20-CE13-0026, MCC, AB); and two SEED FUND ENS LYON-2016&2021 (MCC).

## Author Contributions Statement

**AL**, performed the experiments, analyzed the data, (including statistics), and wrote the paper; **AF**, performed the cloning and the experiment; **AB**, performed the experiment and the quantification **MCC,** performed the experiments and analyzed the data, supervised the work and wrote the paper.

## Data Availability

The *Arabidopsis* lines generated in this study are available from the corresponding author on reasonable request.

---

**Supplemental movie 1.** Time-lapses analysis in Arabidopsis root meristem, expressing Ub10:LifeAct-YFPv (green) together with TUA6-RFP (magenta).

## REFERENCES

Aparicio-Fabre R, Guillen G, Estrada G, Olivares-Grajales J, Gurrola G, Sanchez F (2006) Profilin tyrosine phosphorylation in poly-L-proline-binding regions inhibits binding to phosphoinositide 3-kinase in Phaseolus vulgaris. The Plant journal : for cell and molecular biology 47 (4):491–500. doi:10.1111/j.1365-313X.2006.02787.x

Baluska F, Jasik J, Edelmann HG, Salajova T, Volkmann D (2001) Latrunculin B-induced plant dwarfism: Plant cell elongation is F-actin-dependent. Developmental biology 231 (1):113–124. doi:10.1006/dbio.2000.0115

Buschmann H, Green P, Sambade A, Doonan J, Lloyd C (2011) Cytoskeletal dynamics in interphase, mitosis and cytokinesis analysed through Agrobacterium-mediated transient transformation of tobacco BY-2 cells. New Phytologist 190 (1):258–267

Cho SO, Wick SM (1991) Actin in the developing stomatal complex of winter rye: A comparison of actin antibodies and Rh-phalloidin labeling of control and CB-treated tissues. Cell motility and the cytoskeleton 19 (1):25–36

Clayton L, Lloyd CW (1985) Actin organization during the cell cycle in meristematic plant cells. Actin is present in the cytokinetic phragmoplast. Experimental cell research 156 (1):231–238. doi:10.1016/0014-4827(85)90277-0

Cleary A, Gunning BE, Wasteneys GO, Hepler PK (1992) Microtubule and F-actin dynamics at the division site in living Tradescantia stamen hair cells. Journal of cell science 103 (4):977–988

Dambournet D, Machicoane M, Chesneau L, Sachse M, Rocancourt M, El Marjou A, Formstecher E, Salomon R, Goud B, Echard A (2011) Rab35 GTPase and OCRL phosphatase remodel lipids and F-actin for successful cytokinesis. Nature cell biology 13 (8):981–988. doi:10.1038/ncb2279

Doumane M, Lebecq A, Colin L, Fangain A, Stevens FD, Bareille J, Hamant O, Belkhadir Y, Munnik T, Jaillais Y, Caillaud MC (2021) Inducible depletion of PI(4,5)P_2_ by the synthetic iDePP system in Arabidopsis. Nat Plants 7 (5):587–597. doi:10.1038/s41477-021-00907-z

Doumane M, Lionnet C, Bayle V, Jaillais Y, Caillaud MC (2017) Automated Tracking of Root for Confocal Time-lapse Imaging of Cellular Processes. Bio Protoc 7 (8). doi:10.21769/BioProtoc.2245

Drevensek S, Goussot M, Duroc Y, Christodoulidou A, Steyaert S, Schaefer E, Duvernois E, Grandjean O, Vantard M, Bouchez D, Pastuglia M (2012) The Arabidopsis TRM1-TON1 interaction reveals a recruitment network common to plant cortical microtubule arrays and eukaryotic centrosomes. The Plant cell 24 (1):178–191. doi:10.1105/tpc.111.089748

Durst S, Hedde PN, Brochhausen L, Nick P, Nienhaus GU, Maisch J (2014) Organization of perinuclear actin in live tobacco cells observed by PALM with optical sectioning. Journal of plant physiology 171 (2):97–108. doi:10.1016/j.jplph.2013.10.007

Gujas B, Cruz TMD, Kastanaki E, Vermeer JEM, Munnik T, Rodriguez-Villalon A (2017) Perturbing phosphoinositide homeostasis oppositely affects vascular differentiation in Arabidopsis thaliana roots. Development 144 (19):3578–3589. doi:10.1242/dev.155788

Gungabissoon RA, Jiang C-J, Drøbak BK, Maciver SK, Hussey PJ (1998) Interaction of maize actin-depolymerising factor with actin and phosphoinositides and its inhibition of plant phospholipase C. The Plant Journal 16 (6):689–696. doi:doi:10.1046/j.1365-313x.1998.00339.x

Higaki T, Kutsuna N, Sano T, Hasezawa S (2008) Quantitative analysis of changes in actin microfilament contribution to cell plate development in plant cytokinesis. BMC plant biology 8:80. doi:10.1186/1471-2229-8-80

Hoshino H, Yoneda A, Kumagai F, Hasezawa S (2003) Roles of actin-depleted zone and preprophase band in determining the division site of higher-plant cells, a tobacco BY-2 cell line expressing GFP-tubulin. Protoplasma 222 (3-4):157–165. doi:10.1007/s00709-003-0012-8

Igarashi H, Orii H, Mori H, Shimmen T, Sonobe S (2000) Isolation of a novel 190 kDa protein from tobacco BY-2 cells: possible involvement in the interaction between actin filaments and microtubules. Plant and Cell Physiology 41 (8):920–931

Ischebeck T, Stenzel I, Heilmann I (2008) Type B phosphatidylinositol-4-phosphate 5-kinases mediate Arabidopsis and Nicotiana tabacum pollen tube growth by regulating apical pectin secretion. The Plant cell 20 (12):3312–3330. doi:10.1105/tpc.108.059568

Ischebeck T, Werner S, Krishnamoorthy P, Lerche J, Meijon M, Stenzel I, Lofke C, Wiessner T, Im YJ, Perera IY, Iven T, Feussner I, Busch W, Boss WF, Teichmann T, Hause B, Persson S, Heilmann I (2013) Phosphatidylinositol 4,5-bisphosphate influences PIN polarization by controlling clathrin-mediated membrane trafficking in Arabidopsis. The Plant cell 25 (12):4894–4911. doi:10.1105/tpc.113.116582

Jürgens G (2005) Cytokinesis in higher plants. Annu Rev Plant Biol 56:281–299

Kakimoto T, Shibaoka H (1987) Actin filaments and microtubules in the preprophase band and phragmoplast of tobacco cells. Protoplasma 140 (2-3):151–156

Karimi M, Depicker A, Hilson P (2007) Recombinational cloning with plant gateway vectors. Plant physiology 145 (4):1144–1154. doi:10.1104/pp.107.106989

Katsuta J, Hashiguchi Y, Shibaoka H (1990) The role of the cytoskeleton in positioning of the nucleus in premitotic tobacco BY-2 cells. Journal of cell science 95:413 – 422

Klotz J, Nick P (2012) A novel actin-microtubule cross-linking kinesin, NtKCH, functions in cell expansion and division. The New phytologist 193 (3):576–589. doi:10.1111/j.1469-8137.2011.03944.x

Kojo KH, Higaki T, Kutsuna N, Yoshida Y, Yasuhara H, Hasezawa S (2013) Roles of cortical actin microfilament patterning in division plane orientation in plants. Plant & cell physiology 54 (9):1491–1503. doi:10.1093/pcp/pct093

Kotchoni SO, Zakharova T, Mallery EL, Le J, El-Assal Sel D, Szymanski DB (2009) The association of the Arabidopsis actin-related protein2/3 complex with cell membranes is linked to its assembly status but not its activation. Plant physiology 151 (4):2095–2109. doi:10.1104/pp.109.143859

Kumari P, Dahiya P, Livanos P, Zergiebel L, Kölling M, Poeschl Y, Stamm G, Hermann A, Abel S, Müller S (2021) IQ67 DOMAIN proteins facilitate preprophase band formation and division-plane orientation. Nature Plants 7 (6):739–747

Liao CY, Weijers D (2018) A toolkit for studying cellular reorganization during early embryogenesis in Arabidopsis thaliana. The Plant Journal 93 (6):963–976

Lipka E, Gadeyne A, Stockle D, Zimmermann S, De Jaeger G, Ehrhardt DW, Kirik V, Van Damme D, Muller S (2014) The Phragmoplast-Orienting Kinesin-12 Class Proteins Translate the Positional Information of the Preprophase Band to Establish the Cortical Division Zone in Arabidopsis thaliana. The Plant cell 26 (6):2617–2632. doi:10.1105/tpc.114.124933

Liu B, Palevitz BA (1992) Organization of cortical microfilaments in dividing root cells. Cell motility and the cytoskeleton 23 (4):252–264

Lloyd CW, Traas J (1988) The role of F-actin in determining the division plane of carrot suspension cells. Drug studies. Development 102 (1):211–221

Maeda K, Sasabe M, Hanamata S, Machida Y, Hasezawa S, Higaki T (2020) Actin filament disruption alters phragmoplast microtubule dynamics during the initial phase of plant cytokinesis. Plant and Cell Physiology 61 (3):445–456

Mineyuki Y, Palevitz B (1990) Relationship between preprophase band organization, F-actin and the division site in Allium: Fluorescence and morphometric studies on cytochalasin-treated cells. Journal of cell science 97 (2):283–295

Molchan TM, Valster AH, Hepler PK (2002) Actomyosin promotes cell plate alignment and late lateral expansion in Tradescantia stamen hair cells. Planta 214 (5):683–693

Mole-Bajer J, Bajer AS, Inoue S (1988) Three-dimensional localization and redistribution of F-actin in higher plant mitosis and cell plate formation. Cell motility and the cytoskeleton 10 (1-2):217–228. doi:10.1002/cm.970100126

Muller S, Han S, Smith LG (2006) Two kinesins are involved in the spatial control of cytokinesis in Arabidopsis thaliana. Current biology : CB 16 (9):888–894. doi:10.1016/j.cub.2006.03.034

Nishimura T, Yokota E, Wada T, Shimmen T, Okada K (2003) An Arabidopsis ACT2 dominant-negative mutation, which disturbs F-actin polymerization, reveals its distinctive function in root development. Plant and Cell Physiology 44 (11):1131–1140

Palevitz BA (1987) Actin in the preprophase band of Allium cepa. The Journal of cell biology 104 (6):1515–1519. doi:10.1083/jcb.104.6.1515

Sano T, Higaki T, Oda Y, Hayashi T, Hasezawa S (2005) Appearance of actin microfilament ‘twin peaks’ in mitosis and their function in cell plate formation, as visualized in tobacco BY-2 cells expressing GFP-fimbrin. The Plant journal : for cell and molecular biology 44 (4):595–605. doi:10.1111/j.1365-313X.2005.02558.x

Schaefer E, Belcram K, Uyttewaal M, Duroc Y, Goussot M, Legland D, Laruelle E, de Tauzia-Moreau ML, Pastuglia M, Bouchez D (2017) The preprophase band of microtubules controls the robustness of division orientation in plants. Science 356 (6334):186–189. doi:10.1126/science.aal3016

Schwab B, Mathur J, Saedler R, Schwarz H, Frey B, Scheidegger C, Hülskamp M (2003) Regulation of cell expansion by the DISTORTED genes in Arabidopsis thaliana: actin controls the spatial organization of microtubules. Molecular Genetics and Genomics 269 (3):350–360

Simon ML, Platre MP, Marques-Bueno MM, Armengot L, Stanislas T, Bayle V, Caillaud MC, Jaillais Y (2016) A PtdIns(4)P-driven electrostatic field controls cell membrane identity and signalling in plants. Nat Plants 2:16089. doi:10.1038/nplants.2016.89

Smertenko AP, Deeks MJ, Hussey PJ (2010) Strategies of actin reorganisation in plant cells. Journal of cell science 123 (Pt 17):3019–3028. doi:10.1242/jcs.071126

Smith LG (1999) Divide and conquer: cytokinesis in plant cells. Current opinion in plant biology 2 (6):447–453

Staehelin LA, Hepler PK (1996) Cytokinesis in higher plants. Cell 84 (6):821–824

Staiger CJ, Goodbody KC, Hussey PJ, Valenta R, Drobak BK, Lloyd CW (1993) The profilin multigene family of maize: differential expression of three isoforms. The Plant journal : for cell and molecular biology 4 (4):631–641

Stockle D, Herrmann A, Lipka E, Lauster T, Gavidia R, Zimmermann S, Muller S (2016) Putative RopGAPs impact division plane selection and interact with kinesin-12 POK1. Nat Plants 2:16120. doi:10.1038/nplants.2016.120

Takeuchi M, Karahara I, Kajimura N, Takaoka A, Murata K, Misaki K, Yonemura S, Staehelin LA, Mineyuki Y (2016) Single microfilaments mediate the early steps of microtubule bundling during preprophase band formation in onion cotyledon epidermal cells. Molecular biology of the cell 27 (11):1809–1820. doi:10.1091/mbc.E15-12-0820

Tejos R, Sauer M, Vanneste S, Palacios-Gomez M, Li H, Heilmann M, van Wijk R, Vermeer JE, Heilmann I, Munnik T, Friml J (2014) Bipolar Plasma Membrane Distribution of Phosphoinositides and Their Requirement for Auxin-Mediated Cell Polarity and Patterning in Arabidopsis. The Plant cell 26 (5):2114–2128. doi:10.1105/tpc.114.126185

Voigt B, Timmers AC, Samaj J, Muller J, Baluska F, Menzel D (2005) GFP-FABD2 fusion construct allows in vivo visualization of the dynamic actin cytoskeleton in all cells of Arabidopsis seedlings. European journal of cell biology 84 (6):595–608. doi:10.1016/j.ejcb.2004.11.011

Walker KL, Muller S, Moss D, Ehrhardt DW, Smith LG (2007) Arabidopsis TANGLED identifies the division plane throughout mitosis and cytokinesis. Current biology : CB 17 (21):1827–1836. doi:10.1016/j.cub.2007.09.063

Wu SZ, Bezanilla M (2014) Myosin VIII associates with microtubule ends and together with actin plays a role in guiding plant cell division. Elife 3. doi:10.7554/eLife.03498

Xu XM, Zhao Q, Rodrigo-Peiris T, Brkljacic J, He CS, Muller S, Meier I (2008) RanGAP1 is a continuous marker of the Arabidopsis cell division plane. Proceedings of the National Academy of Sciences of the United States of America 105 (47):18637–18642. doi:10.1073/pnas.0806157105

Yoneda A, Akatsuka M, Kumagai F, Hasezawa S (2004) Disruption of actin microfilaments causes cortical microtubule disorganization and extra-phragmoplast formation at M/G1 interface in synchronized tobacco cells. Plant & cell physiology 45 (6):761–769. doi:10.1093/pcp/pch091

